# New and improved GRAB fluorescent sensors for monitoring dopaminergic activity *in vivo*

**DOI:** 10.1101/2020.03.28.013722

**Authors:** Fangmiao Sun, Jingheng Zhou, Bing Dai, Tongrui Qian, Jianzhi Zeng, Xuelin Li, Yizhou Zhuo, Yajun Zhang, Ke Tan, Jiesi Feng, Hui Dong, Cheng Qian, Dayu Lin, Guohong Cui, Yulong Li

**Affiliations:** State Key Laboratory of Membrane Biology, Peking University School of Life Sciences, 100871 Beijing, China; PKU-IDG/McGovern Institute for Brain Research, 100871 Beijing, China; Peking-Tsinghua Center for Life Sciences, 100871 Beijing, China; School of Life Sciences, Tsinghua University, Beijing 100084, China; Neurobiology Laboratory, National Institute of Environmental Health Sciences, National Institutes of Health, Research Triangle Park, NC 27709, USA; Neuroscience Institute, Department of Psychiatry, New York University School of Medicine, New York, NY 10016, USA

## Abstract

The monoamine neuromodulator dopamine (DA) plays a critical role in the brain, and the ability to directly measure dopaminergic activity is essential for understanding its physiological functions. We therefore developed the first red fluorescent GPCR-activation–based DA (GRAB_DA_) sensors and optimized versions of green fluorescent GRAB_DA_ sensors following our previous studies. In response to extracellular DA, both the red and green GRAB_DA_ sensors have a large increase in fluorescence (ΔF/F_0_ values of 150% and 340%, respectively), with subcellular resolution, subsecond kinetics, and nanomolar to submicromolar affinity. Moreover, both the red and green GRAB_DA_ sensors readily resolve evoked DA release in mouse brain slices, detect compartmental DA release in live flies with single-cell resolution, and report optogenetically elicited nigrostriatal DA release as well as mesoaccumbens dopaminergic activity during sexual behavior in freely behaving mice. Importantly, co-expressing red GRAB_DA_ with either green GRAB_DA_ or the calcium indicator GCaMP6s provides a robust tool for simultaneously tracking neuronal activity and dopaminergic signaling in distinct circuits *in vivo*.

---

Dopamine (DA) is an essential monoamine neuromodulator produced primarily in the midbrain and released throughout the central nervous system. A multitude of brain functions are regulated by DA, including motor control, motivation, learning and memory, and emotional control^1–9^. Consistent with these key physiological roles, altered DA signaling has been implicated in a variety of brain disorders, including Parkinson’s disease, addiction, schizophrenia, attention-deficit/hyperactivity disorder, and posttraumatic stress disorder^10–20^. Thus, tools that can sense changes in DA concentration with high spatiotemporal resolution, high specificity, and high sensitivity will greatly facilitate our study of the diverse functions that the dopaminergic system plays under both physiological and pathological conditions.

Previous techniques for measuring DA dynamics, including microdialysis, electrochemical probes, reporter cells, and gene expression–based assays, lack sufficient spatiotemporal resolution and/or molecular specificity^21–30^. Recently, our group^31^ and Patriarchi et al.^32^ independently developed two series of genetically encoded, G-protein–coupled receptor (GPCR)–based DA sensors called GRAB_DA_ and dLight, respectively. Taking advantage of naturally occurring DA receptors, these sensors convert a ligand–stabilized conformational change in the DA receptor into an optical response via a conformation-sensitive fluorescent protein inserted in the receptor’s third intracellular loop. Our first-generation DA receptor–based sensors called GRABd_DA1m_ and GRAB_DA1h_ were used to detect cell type–specific DA dynamics in several organisms, including *Drosophila,* zebrafish, mice, and zebra finches^31,33–35^. Here, we employed semi-rational engineering to modify the green fluorescent protein. The resulted second-generation sensors called GRAB_DA2m_ and GRAB_DA2h_ (abbreviated DA2m and DA2h, respectively) have a 2–3-fold higher dynamic range and improved *in vivo* performance in comparison to their corresponding first-generation sensors.

Red fluorescent sensors have distinct and well-separated spectra from those of GFP–based sensors and blue light–excitable channelrhodopsin 2 (ChR2), thus enabling the orthogonal readout of two neurochemical events or simultaneous monitoring blue light–mediated activity control. Moreover, red fluorescent sensors require an excitation light with a relatively longer wavelength, providing additional advantages over green fluorescent proteins, including reduced phototoxicity, reduced background, and deeper tissue penetration^36,37^. Starting with the dopamine D2 receptor (D_2_R) and a conformation-sensitive red fluorescent protein cpmApple^38^, we generated a series of red fluorescent DA sensors called rGRAB_DA1m_ and rGRAB_DA1h_ (abbreviated rDA1m and rDA1h, respectively), with a dynamic range similar to the corresponding first-generation green DA sensors.

Here, we report the development, *in vitro* characterization, and *in vivo* application of our novel red fluorescent DA sensors and second-generation green fluorescent DA sensors for monitoring DA dynamics in real-time in *Drosophila* and mice in response to physiologically relevant stimuli and during complex behaviors.

## Results

### Development and *in vitro* characterization of DA sensors

To develop red fluorescent DA sensors, we systematically optimized the cpmApple insertion sites within the third intracellular loop of D_2_R, the linker sequences, and the cpmApple module itself, using both brightness and DA–induced change in fluorescence (ΔF/F_0_) as our selection criteria. Screening a library containing over 2000 sensor variants revealed the sensor with the highest fluorescence response; we called this sensor rDA0.5. Next, we used a rational strategy to introduce an iterative series of mutations in the D_2_R module of rDA0.5, generating versions with differing apparent affinities to DA. The medium-affinity sensor rDA1m was generated by introducing the K367^6.29^L mutation in the receptor, and the high-affinity sensor rDA1h was generated by introducing the T205^5.54^M mutation in rDA1m. Finally a DA–insensitive version (rDA-mut) was generated by introducing the C118^3.36^A and S193^5.42^N mutations in rDA1h (Fig. 1a,b,d and Supplementary Fig. S1b,c). All three versions localized nicely on the cell membrane when expressed in HEK293T cells (Fig. 1c). rDA1m and rDA1h had an EC_50_ of 95 nM and 4 nM, respectively (Fig. 1d). Moreover, the application of 100 μM DA elicited a 150% and 100% increase in fluorescence of rDA1m and rDA1h, respectively, which was blocked by the D_2_R antagonist haloperidol (Halo) (Fig. 1c,d). As expected, DA had no effect in cells expressing rDA-mut, even at the highest concentration tested (Fig. 1c,d).

**Fig. 1.**
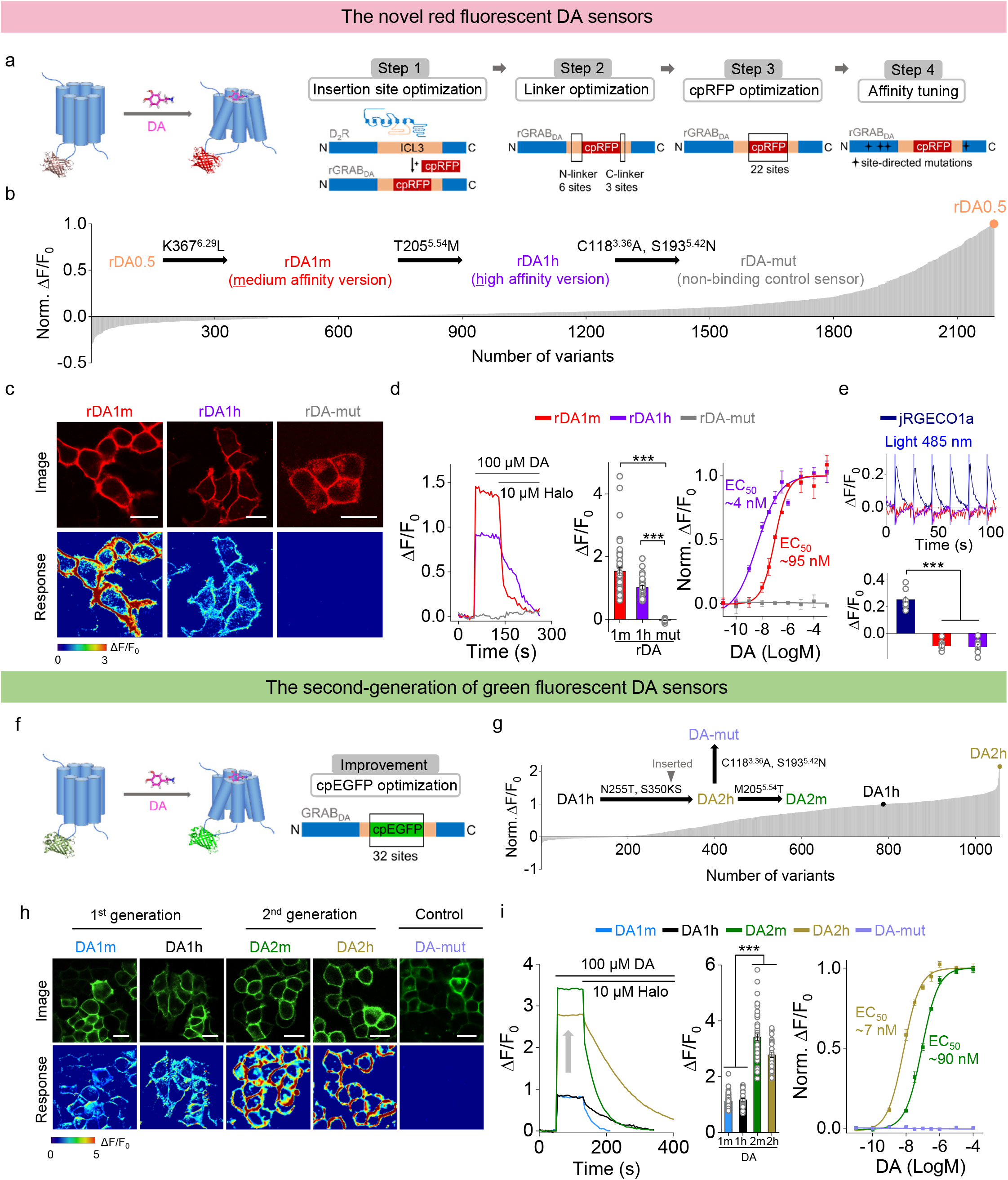
Development and characterization of two novel red fluorescent DA sensors and second-generation green fluorescent DA sensors. **a, f**, Schematic illustration showing the design and optimization of the red and green fluorescent GRAB_DA_ sensors. **b**, Normalized change in fluorescence in response to 100 μM DA measured for red fluorescent DA sensor variants during steps 1–3. A total of 2189 variants were screened, and the variant with the highest fluorescence change (named rDA0.5) was selected for subsequent affinity tuning; rDA0.5 was then sequentially mutated as shown to generate rDA1m, rDA1h, and rDA-mut. **c**, Representative red fluorescence images (top) and change in fluorescence in response to 100 μM DA (bottom) in HEK293T cells expressing rDA1m, rDA1h or rDA-mut. Scale bars, 20 μm. **d**, Representative traces (left), group summary of peak ΔF/F_0_ (middle; n=17–46 cells), and normalized dose-response curves (right; n=3 wells with 200–400 cells/well) in response to DA. Where indicated, 10 μM haloperidol (Halo) was applied. **e**, Representative traces (top) and group summary (bottom; n=8–9 cells) of ΔF/F_0_ in response to blue light in cells expressing jRGECO1a, rDA1m, or rDA1h. **g**, Normalized ΔF/F_0_ in response to 100 μM DA application for 1056 variants, normalized to the first-generation DA1h sensor. DA2h was then mutated as shown to generate DA2m and DA-mut. **h**, Representative green fluorescence images (top) and change in fluorescence in response to 100 μM DA (bottom) in HEK293T cells expressing DA1m, DA1h, DA2m, DA2h, or DA-mut. Scale bars, 20 μm. **i**, Representative traces (left), group summary of peak ΔF/F_0_ (middle; n=33–68 cells), and normalized dose-response curves (right; n=3 wells with 200–500 cells/well) in response to DA. Where indicated, 10 μM Halo was applied.

Next, we asked whether the red DA sensors undergo photoactivation when expressed in HEK293T cells and cultured neurons. Previous studies showed that cpmApple–based sensors can undergo photoactivation when illuminated with blue light^39–41^, preventing the combined use of these sensors with ChR2. We found that although cpmApple–based red fluorescent calcium indicator jRGECO1a^41^ had a 20% increase in fluorescence upon blue light illumination (Fig. 1e), blue light had no effect on the fluorescence of either rDA1m or rDA1h, suggesting that our red fluorescent DA sensors are compatible for use with blue light–mediated optogenetic activation. Moreover, we found that photostability of the red DA sensors was similar to or better than several commonly used red fluorescent proteins (Supplementary Fig. S1d).

In parallel, we optimized our first-generation green fluorescent DA sensors by performing random mutagenesis at 32 sites in the cpEGFP module (Fig. 1f). Screening ∼1000 variants yielded DA2h, the second-generation high-affinity green fluorescent DA sensor (Fig. 1g and Supplementary Fig. S1a); this sensor was then used to generate the medium-affinity DA2m and DA–insensitive DA-mut versions (Fig. 1g). Compared to their corresponding first-generation DA1m and DA1h sensors, the second-generation green fluorescent DA sensors had a 2–3 fold higher fluorescence increase (ΔF/F_0_ ∼340% for DA2m and ∼280% for DA2h) to 100 μM DA while maintaining their apparent affinity to DA, with EC_50_ values of 90 nM and 7 nM for DA2m and DA2h, respectively (Fig. 1h,i). Finally, the DA–insensitive DA-mut sensor localized well on the cell membrane but did not respond to DA (Fig. 1h,i).

We further characterized the specificity, kinetics, and downstream coupling of our newly developed DA sensors. With respect to specificity, the DA–induced signals of both the red and green DA sensors were blocked by the D_2_R–specific antagonists Halo and eticlopride, but not the D_1_R antagonist SCH-23390. None of these four DA sensors responded to a variety of neurotransmitters and neuromodulators (Fig. 2a). We also found that although norepinephrine (NE) is structurally similar to DA, both the red and green DA sensors were 10–20 fold more selective for DA over NE (Fig. 2a, inset), suggesting the high selectivity of the sensors to DA at physiologically relevant concentrations^42–45^.

**Fig. 2.**
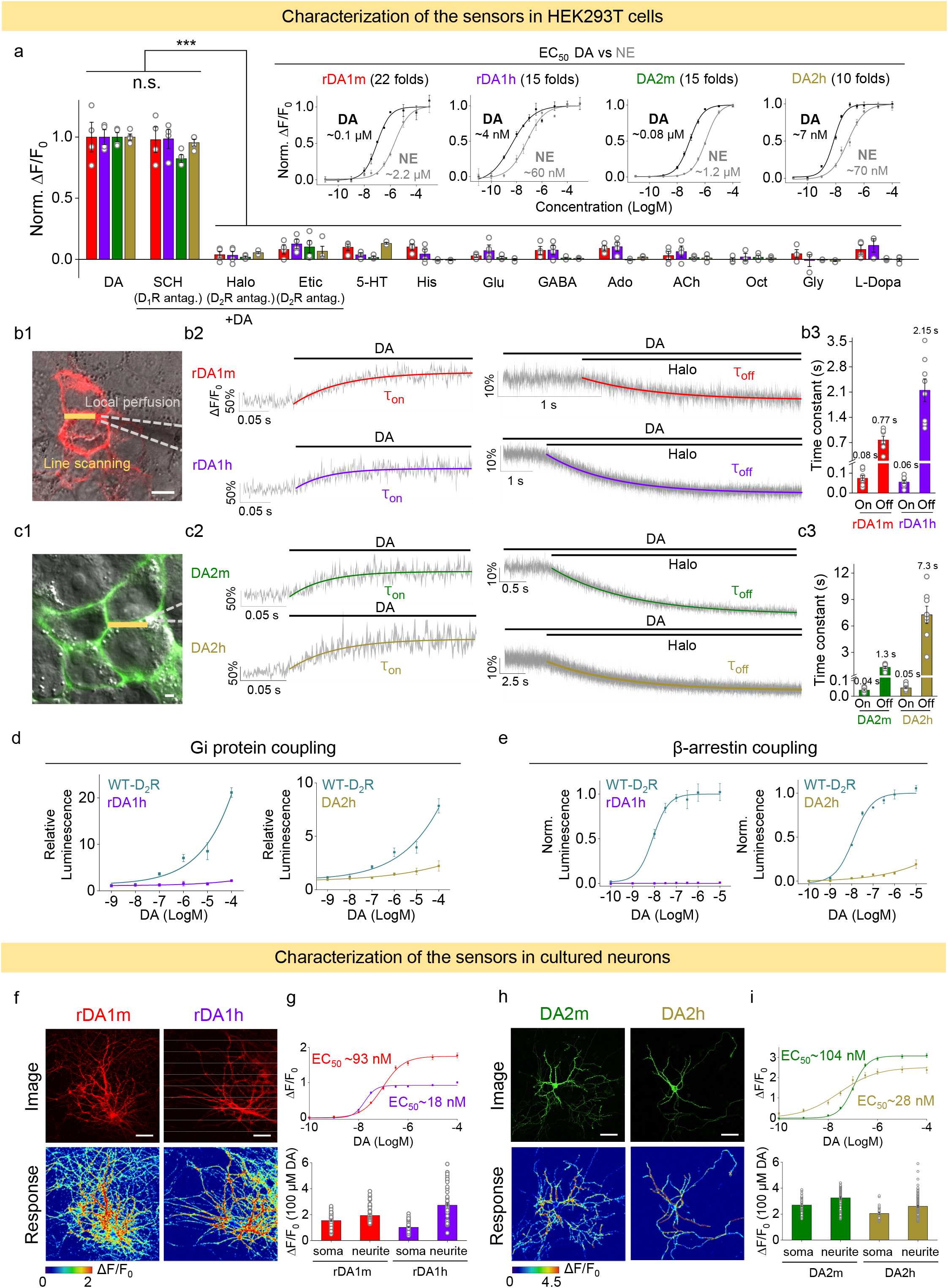
Characterization of GRAB_DA_ sensors in HEK293T cells and cultured neurons. **a**, Normalized ΔF/F_0_ was measured in HEK293T cells expressing rDA1m, rDA1h, DA2m, or DA2h following the application of DA alone, DA+SCH-23390, DA+haloperidol, DA+eticlopride, serotonin (5-HT), histamine, glutamate, gamma-aminobutyric acid (GABA), adenosine, acetylcholine (ACh), octopamine, glycine, or L-DOPA (all applied at 1 μM); n=3–4 wells with 200–1200 cells/well. The insets show normalized dose-response curves for DA and norepinephrine (NE) in cells expressing rDA1m, rDA1h, DA2m, or DA2h (n=3 wells with 200–800 cells/well). **b-c**, Response kinetics for rDA1m, rDA1h, DA2m, and DA2h. **b1** and **c1**, Schematic illustration showing the local perfusion system for applying compounds. DA and Halo were delivered via a glass pipette (dashed gray lines) positioned at the sensor-expressing cells, and the fluorescence signal was measured using confocal line scanning (thick yellow lines). Scale bars, 10 μm. **b2** and **c2**, Representative traces showing fluorescence after application of DA (left) and subsequent addition of Halo (right). The traces were the average of 3 different regions of interest (ROIs) on the scanning line, shaded with ± s.e.m.. Each trace was fitted with a single-exponential function to determine τ_on_ (left) and τ_off_ (right). **b3** and **c3**, Group summary of τ_on_ and τ_off_ (n=7–10 cells). **d**, Gi coupling of wild-type (WT) D_2_R, rDA1h, and DA2h was measured using the luciferase complementation assay (n=3 wells each). **e**, β-arrestin coupling of WT D_2_R, rDA1h, and DA2h was measured using the TANGO assay (n=3 wells each). **f, h**, Representative red (**f**) and green (**h**) fluorescence images (top) and response to 100 μM DA (bottom) in neurons expressing the indicated sensors. Scale bars, 10 μm. **g,i**, Dose-response curves (top) and group summary (bottom) of the responses measured in the soma and neurites of neurons expressing the indicated sensors (n=21–60 neurons).

We next characterized the kinetics of the DA sensors using rapid line scanning in response to a local puff of DA (to measure τ_on_) followed by Halo (to measure τ_off_) to sensor-expressing HEK293T cells. As shown in Fig. 2b and 2c, τ_on_ was <100 ms for all four DA sensors while the high-affinity versions of the red and green fluorescent sensors had relatively slower off kinetics compared to their corresponding medium-affinity counterparts.

To examine whether our DA sensors couple to downstream signaling pathways, we used the luciferase complementation assay^46^ and TANGO assay^47^ to measure activation of the Gi (Fig. 2d) and β-arrestin (Fig. 2e) pathways, respectively. When expressed in HEK293T cells, both the rDA1h and DA2h sensors had virtually no downstream coupling; as a control, the wild-type D_2_R showed robust coupling (Fig. 2d,e). In the cultured neurons, we also found that both the red and green DA sensors readily localized on the cell membrane and responded well to DA (Fig. 2f-i), supporting the utility of the sensors in the nervous system.

Lastly, we compared the properties of the second-generation green fluorescent DA2m sensor with D_1_R-based dLight1.1 and dLight 1.2 sensors (Supplementary Fig. S2a-j). When expressed in cultured cells, DA2m had a higher apparent affinity to DA, higher basal brightness, a higher ΔF/F_0_ response, and a higher signal-to-noise ratio (SNR) than dLight.

Taken together, these results indicate that our red and improved green fluorescent DA sensors are highly sensitive and specific to DA, with rapid kinetics, suggesting that they will be suitable for monitoring dopaminergic activity *in vivo*.

### Imaging of DA release in acute mouse brain slices

Next, we examined whether our DA sensors can be used to measure the release of endogenous DA in acute brain slices. We first injected AAVs expressing rDA1m, rDA1h, or DA2m into the nucleus accumbens (NAc), which receives strong innervation from midbrain dopaminergic neurons^48–50^ (Fig. 3a,b). Two weeks after expressing the sensor, we prepared acute brain slices and used 2-photon imaging combined with electrical stimulation to measure stimulus–evoked DA release (Fig. 3a). Electrical stimuli delivered at 20 Hz induced progressively increased fluorescence responses in the NAc, which was blocked by Halo (Fig. 3c,e). We also measured the sensors’ kinetics during 10 pulses applied at 100 Hz. The τ_on_ and τ_off_ values are 0.08–0.15 s and 5.2–11.8 s, respectively (Fig. 3d).

**Fig. 3.**
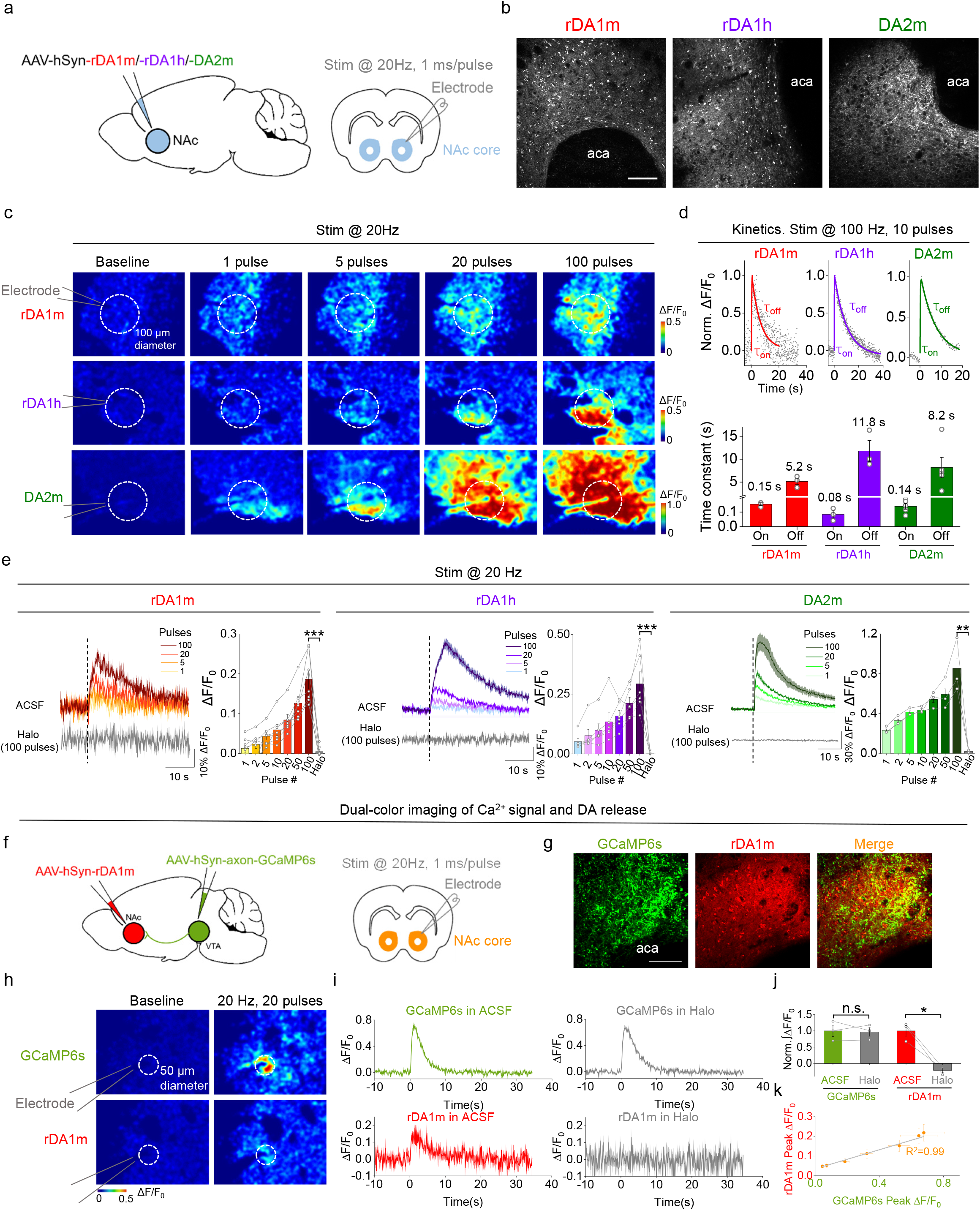
GRAB_DA_ sensors can be used to measure DA release in acute mouse brain slices. **a**, Schematic illustration depicting the experimental design for injecting AAVs expressing the rDA1m, rDA1h, or DA2m sensors in the mouse NAc, followed by imaging of electrical stimulation-evoked DA release in the NAc in acute brain slices. **b**, Representative fluorescence images showing the expression of rDA1m, rDA1h, and DA2m in the NAc. The anterior cingulate area (aca) adjacent to the NAc is indicated. Scale bar, 100 μm. **c**, Fluorescence images measured in brain slices from mice expressing rDA1m, rDA1h, or DA2m in the NAc. The dashed circles indicate the ROIs used to analyze the signals. **d**, Representative traces showing the normalized change in fluorescence (top) and group summary of τ_on_ and τ_off_ (bottom) for rDA1m, rDA1h, and DA2m in response to 10 electrical stimuli applied at 100 Hz (n=3–5 slices from 2–3 mice). The data were processed with 2× binning. **e**, Representative traces and group summary of the change in fluorescence measured for rDA1m (left), rDA1h (middle), and DA2m (right) in response to the indicated number of electrical stimuli applied at 20 Hz; where indicated, Halo (10 μM) was included in the bath solution (n=3–7 slices from 2–4 mice). **f**, Schematic illustration depicting the strategy for injecting AAVs in the NAc and VTA, followed by simultaneous recording of Ca^2+^ and DA release using dual-color imaging. **g**, Representative green fluorescence (GCaMP6s), red fluorescence (rDA1m), and merged images of the NAc. The aca adjacent to the NAc is indicated. Scale bar, 100 μm. **h**, Fluorescence images showing the response of axon-GCaMP6s and rDA1m following 20 electrical stimuli applied at 20 Hz. The dashed circles indicate the ROIs used to analyze the signals. **i-j**, Representative traces (**i**) and group summary (**j**) of the change in axon-GCaMP6s and rDA1m fluorescence in response to 20 electrical stimuli applied at 20 Hz in the absence or presence of Halo (10 μM); n=3 slices from 3 mice. **k**, The peak change in rDA1m fluorescence plotted against the peak change in axon-GCaMP6s fluorescence in response to various numbers of pulses applied at 20 Hz. The data were fitted to a linear function, and the coefficient of correlation is shown (n=8 slices from 3 mice). Average traces shaded with ± s.e.m from one slice are shown for representation.

To test whether the red fluorescent DA sensor is spectrally compatible with the green fluorescent calcium sensor, we co-expressed the axon–targeted GCaMP6s^51^ in the ventral tegmental area (VTA) and rDA1m in the NAc, then simultaneously imaged calcium and DA in the NAc during 20 Hz electrical stimulation (Fig. 3f,g). We found that the electrical stimulation evoked robust fluorescence increase of both GCaMP6s and rDA1m and their magnitudes of increases were highly correlated (Fig. 3k). Application of the D_2_R antagonist Halo blocked the rDA1m response but had no effect on the GCaMP6s response (Fig. 3h-j). Taken together, these data indicate that the rDA1m, rDA1h, and DA2m sensors can detect dopaminergic activity in brain slices with high specificity, sensitivity and temporospatial resolution. Moreover, our results confirm that the red fluorescent DA sensors are spectrally compatible with green fluorescent probes, allowing for simultaneous dual-color imaging.

### *In vivo* imaging of DA in *Drosophila*

In *Drosophila,* dopaminergic activity in the mushroom body (MB) has been found to be both necessary and sufficient for the associate learning between odor and aversive experience, e.g. body shock^52–55^. Next, we generated a transgenic *Drosophila* expressing rDA1m in Kenyon cells (KCs) in the olfactory MB and measured the fluorescence level of the rDA1m sensor using *in vivo* 2-photon imaging when physiologically relevant stimuli were presented (Fig. 4a,b). When we delivered either the odorant or body shock, we observed a time–locked fluorescence increase in the medial lobe of the MB; this increase was blocked by pretreating the animals with Halo (Fig. 4c,d) and was not observed in flies expressing the DA–insensitive rDA-mut sensor. Importantly, the endogenous signal did not saturate the rDA1m sensor’s response, as application of 100 μM DA caused a significantly larger, sustained increase in fluorescence (Fig. 4e).

**Fig. 4.**
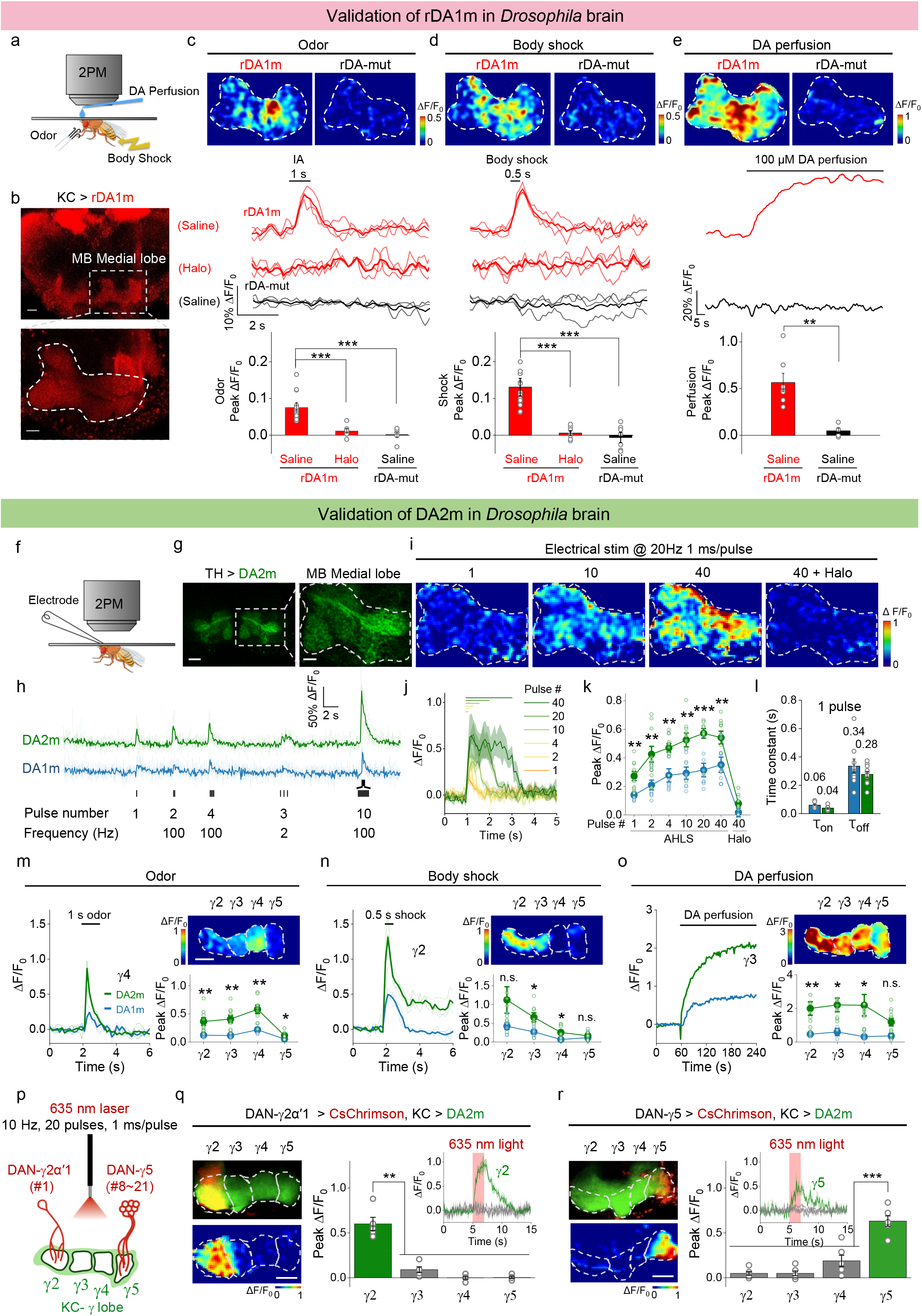
*In vivo* 2-photon imaging of DA dynamics in *Drosophila* using GRAB_DA_ sensors. **a**, Schematic illustration depicting the experimental setup for imaging fluorescence changes in *Drosophila* expressing GRAB_DA_ sensors using 2-photon microscopy (2PM). Application of various stimuli are also depicted. **b**, Representative red fluorescence images of a transgenic fly expressing rDA1m in the Kenyon cells (KCs), with an expanded view of the olfactory mushroom body (MB) medial lobe. Scale bars, 20 μm (top) and 10 μm (bottom). **c-e**, Representative images (top; the dashed area indicates the MB medial lobe), traces (middle), and group summary (bottom) of ΔF/F_0_ in response to a 1 s application of odorant (**c**), a 500 ms body shock (**d**), and 100 μM DA (**e**) in flies expressing rDA1m in saline, rDA1m in Halo, or rDA1m-mut in saline (n=5–15 flies each). **f**, Schematic illustration depicting the experimental setup for imaging electrical stimulation–evoked DA release. **g**, Representative green fluorescence images of a transgenic fly expressing DA2m in dopaminergic neurons (DANs), with an expanded view of the MB medial lobe. Scale bars, 20 μm (left) and 10 μm (right). **h**, Representative traces of DA2m and DA1m fluorescence; where indicated, electrical stimuli were applied. **i, j**, Representative images (**i**, the dashed area indicates the MB medial lobe) and traces (**j**) of DA2m ΔF/F_0_ in response to electrical stimuli applied at 20 Hz in the absence or presence of Halo. **k**, Group summary of DA2m and DA1m ΔF/F_0_ in response to electrical stimuli applied at 20 Hz in the absence or presence of Halo (n=9–10 flies each). **l**, Kinetics (τ_on_ and τ_off_) of DA2m and DA1m in response to a single electrical stimulus (n=9–10 flies each). **m-o**, Representative traces (left), fluorescence images (upper right), and group summary (bottom right) of the indicated MB lobe compartments in response to a 1 s application of odorant (**m**), a 500 ms body shock (**n**), and 100 μM DA (**o**); n=3-10 flies each. **p**, Schematic illustration depicting the strategy for imaging optogenetically induced DA release. DA2m is expressed in the KCs, and CsChrimson is expressed in the DANs in either the γ2 or γ5 MB compartment (with the number of innervating cells indicated). **q, r**, Representative fluorescence images of CsChrimson (upper left), DA2m (lower left), representative traces (top right), and group summary (bottom right) of DA2m fluorescence in the γ2 (**q**) and γ5 (**r**) MB compartments in response to optogenetic stimulation (n=5–6 flies each). Scale bars, 20 μm. The group data for DA1m shown in panels **k, m, n**, and **o** were reproduced from Sun et al.^31^ with permission. Average traces (bold), overlaid with single-trial traces (light) or shaded with ± s.e.m, from one fly are shown for representation.

We then compared *in vivo* performance between the first-generation and second-generation green fluorescent DA sensors in flies. Electrical stimulation of the MB medial lobe elicited robust fluorescence increase in nearby DA sensor-expressing dopaminergic neurons (DANs) with a higher response observed in DA2m-expressing flies (Fig. 4f,g). The temporal dynamics of the DA2m and DA1m responses (τ_on_ and τ_off_) were similar (Fig. 4l) and both responses were blocked by Halo (Fig. 4h-k). The spatial patterns of the responses in the MB during odorant or body shock delivery were also similar between DA2m- and DA1m-expressing flies^31^ while consistently higher responses were observed in DA2m-expressing flies (Fig. 4m-o). In a separate experiment, we compared the *in vivo* performance of DA2m with dLight1.3b, which has the highest dynamic range among the dLight series of sensors, and found that DA2m produced a 3-fold larger response (Supplementary Fig. S2k-o).

Previous studies have found that the *Drosophila* MB is innervated by the axonal neuropils from 15 DAN subgroups, with each lobe containing axons from one to approximately two dozens of neurons^56,57,58^. We next asked whether our highly sensitive DA2m sensor could detect DA release in individual MB compartments, even when the source of DA is from a single neuron. As a proof-of-principle experiment, we co-expressed the red light–activated channelrhodopsin CsChrimson^59^ in DANs and DA2m in KCs. We then used 635 nm wavelength light to activate the only DAN neuron innervating the γ2α’1 compartment or the 8–21 neurons innervating the γ5 compartment while measuring DA2m fluorescence (Fig. 4p). We found that the change in DA2m fluorescence was both time–locked to the CsChrimson activation and spatially confined to the respective compartments, supporting functional independence among MB compartments (Fig. 4q,r).

To confirm that the DA2m sensor does not couple to Gi signaling *in vivo,* we used cAMP level as a proxy for Gi signaling and expressed the red fluorescent cAMP sensor Pink-Flamindo^60^ to measure body shock–evoked increase in cAMP in KCs. We found that the presence of the DA2m sensor had no effect on cAMP production (Supplementary Fig. S3). Thus, these data confirm that the DA2m sensor does not affect the endogenous signaling pathways *in vivo*.

### Red and green fluorescent DA sensors can be used to detect optogenetically induced nigrostriatal DA release in freely moving mice

To demonstrate that our new DA sensors can be used to measure dopaminergic activity in freely moving animals, we measured DA dynamics in the mouse dorsal striatum, which receives motor control–related nigrostriatal dopaminergic projections from the substantia nigra pars compacta (SNc)^61^, by expressing the optogenetic tool C1V1^62^ in the SNc and various DA sensors in the dorsal striatum. To provide a frame of reference for the red and green fluorescent sensors, we also co-expressed EGFP or tdTomato, respectively, in the dorsal striatum (Fig. 5a,b). C1V1–mediated optogenetic activation of DANs in the SNc elicited a robust transient increase in DA sensor fluorescence in the dorsal striatum (Fig. 5e,g,i,k). Importantly, the signal was prolonged by the DA transporter blocker methylphenidate and blocked by the D_2_R antagonist eticlopride (Fig. 5e-l); in contrast, the signal was unaffected by the NE transporter blocker desipramine and the α2-adrenergic receptor antagonist yohimbine (Supplementary Fig. S4). As a control, the DA–insensitive sensor rDA-mut did not respond to C1V1–mediated optogenetic activation (Fig. 5c,d). Finally, both EGFP fluorescence and tdTomato fluorescence were unchanged throughout the experiments, indicating a lack of movement–related artifacts during recording. Thus, our new DA sensors can detect optogenetically induced DA release in freely moving mice with high sensitivity and high molecular specificity.

**Fig. 5.**
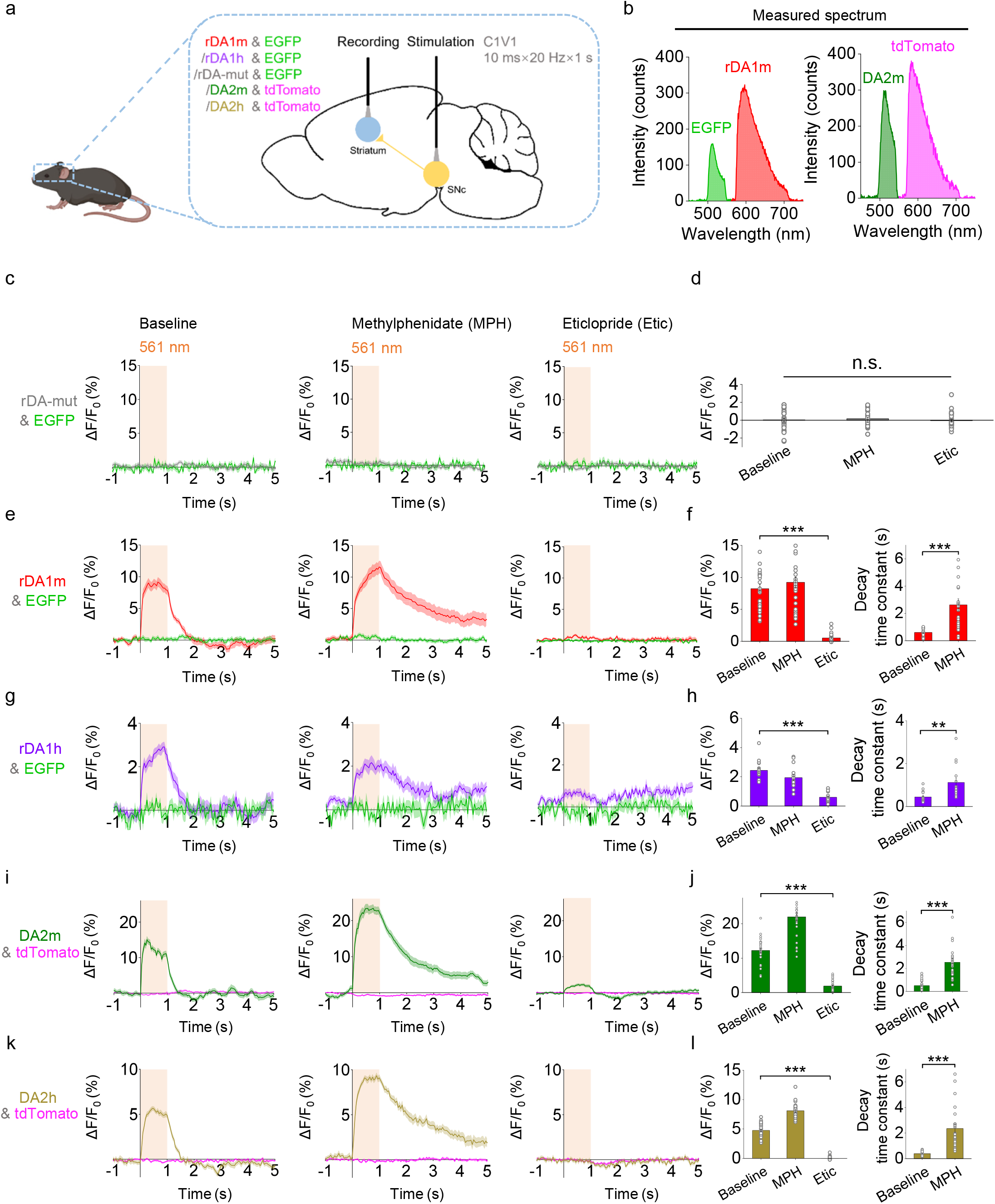
GRAB_DA_ sensors can detect optogenetically induced nigrostriatal DA release in freely moving mice. **a**, Schematic illustration depicting the experimental setup for dual-color fiber photometry recording in the dorsal striatum while optogenetically stimulating DANs in the SNc. The combinations of sensor and fluorescent protein were used in the experiments shown in panels **c-l**. **b**, Measured emission spectra of rDA1m and EGFP (left) and DA2m and tdTomato (right). **c, e,g,i,k**, Average ΔF/F_0_ traces of the indicated sensors and fluorescent proteins during optogenetic stimulation under control conditions (left) or in the presence of methylphenidate (MPH) or eticlopride (Etic). **d, f,h,j,l**, Group summary of ΔF/F_0_ and τ_off_ (where applicable) for the corresponding sensors in panels **c, e, g, i**, and **k**, respectively (n=15–30 trials from 3–6 hemispheres of 3–4 mice per condition).

### Detection of dopaminergic activity in the mouse NAc during sexual behavior

We previously showed^31^ that dopaminergic activity can be measured during male sexual behavior^4^. To ask whether our improved DA2h performs better than the DA1h sensor in reporting real-time DA dynamics in freely moving mice, we expressed DA1h in one side of the NAc and DA2h in the contralateral NAc, and performed bilateral fiber photometry recording during mating (Fig. 6a,b). We found that the DA1h and DA2h signals were closely correlated in time. Consistent with the improved performance of DA2h in other systems, the DA2h sensor had a significantly higher fluorescence change (ΔF/F_0_) than DA1h during various stages of sexual behavior (Fig. 6c-f).

**Fig. 6.**
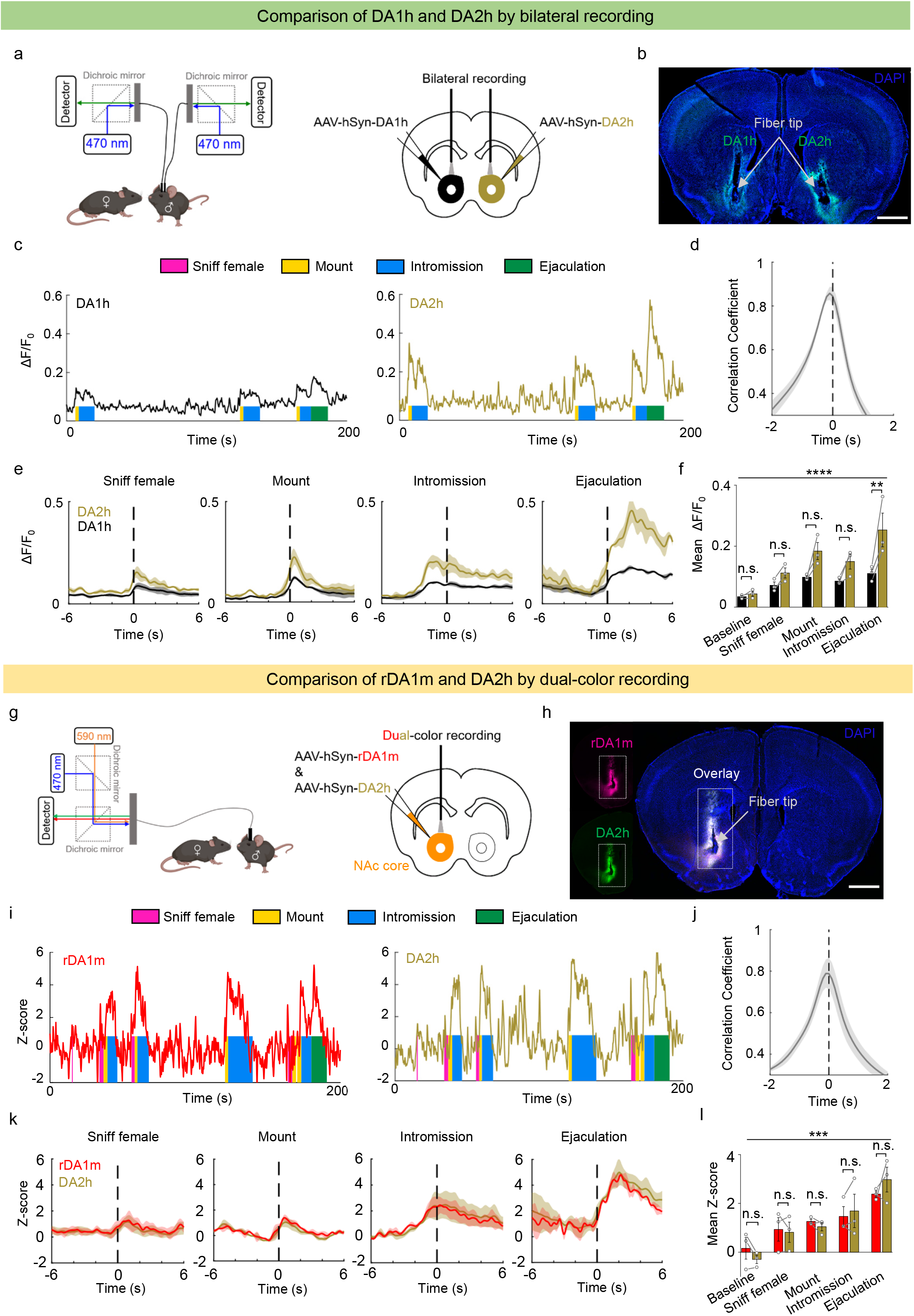
GRAB_DA_ sensors can be used to measure dopaminergic activity in the mouse NAc during sexual behavior. **a**, Schematic illustration depicting the strategy for injecting AAVs and performing bilateral fiber photometry recording. **b**, Representative image showing the expression of DA1h and DA2h in opposite hemispheres, as well as the recording site (‘fiber tip’). The nuclei were counterstained with DAPI (blue). Scale bar, 1 mm. **c**, Representative traces of DA1h and DA2h ΔF/F_0_ measured during the indicated stages of mating. **d**, Summary of the time shift correlation coefficient between the DA1h and DA2h signals (n=3 mice). **e**, Average post-stimulus histograms showing the ΔF/F_0_ signals of DA1h and DA2h aligned to the onset of the indicated mating events (n=3 mice). **f**, Group summary of mean ΔF/F_0_ measured for DA1h and DA2h during the indicated mating events (n=3 mice). **g**, Schematic illustration depicting the strategy for injecting AAVs and performing dual-color fiber photometry recording. **h**, Representative images showing the colocalized expression of rDA1m (red) and DA2h (green), as well as the recording site. The nuclei were counterstained with DAPI (blue). Scale bar, 1 mm. **i**, Representative traces of the concurrent *Z*-score signals of rDA1m (left) and DA2h (right) during the indicated stages of sexual behavior. **j**, Summary of the time shift correlation coefficient between the rDA1m and DA2h signals (n=3 mice). **k**, Average post-stimulus histograms showing the *Z*-score signals of rDA1m and DA2h aligned to the onset of the indicated mating events (n=3 mice). **l**, Group summary of the mean *Z*-scores measured for rDA1m and DA2h during the indicated mating events (n=3 mice).

We next compared the performance of red fluorescent rDA1m sensor with that of the best green fluorescent DA sensor, DA2h. We co-injected viruses expressing rDA1m and DA2h into the NAc core and performed dual-color fiber photometry recording three weeks after virus injection (Fig. 6g,h). We found that despite the dynamic range (ΔF/F_0_) of the rDA1m is smaller than that of DA2h (Supplementary Fig. S5 and S6), rDA1m and DA2h detected qualitatively similar DA release during sexual behavior when the responses are Z scores (Fig. 6i,k,l). The moment-to-moment correlation coefficient between rDA1m and DA2h is similar to that between DA1h and DA2h (Fig. 6d,j). Importantly, we did not observe crosstalk between the red and green DA sensors as no signal in the red channel was detected when only 470 nm light was delivered and *vice versa* (Supplementary Fig. S6). Taken together, the rDA1m is capable of detecting DA release *in vivo* during natural behaviors and the behavior–related DA responses detected by the red and green DA sensors are qualitatively similar.

## Discussion

Here, we report the development and characterization of a new set of genetically encoded DA sensors. Moreover, we show that these sensors can be used to measure DA release in mouse brain slices, as well as in the *Drosophila* olfactory system and the NAc in freely moving mice during sexual behavior. Importantly, these sensors can report DA release evoked by electrical stimulation, optogenetic activation, and various physiologically relevant stimuli and behaviors. The availability of both high-affinity and medium-affinity versions provides the opportunity to probe DA dynamics over a broad range of concentrations. Moreover, the DA2m sensor provides an ideal balance of sensitivity, dynamic range, and brightness, making it a powerful tool for monitoring DA dynamics *in vivo* with a high signal-to-noise ratio in several models, including flies and mice^63,64^.

DA has long been regarded as a neuromodulator that can diffuse broadly and exert its effects over a large volume with both slow and fast kinetics as well as tonic and phasic release patterns. However, several key questions regarding dopaminergic activity remain unanswered. For example, how are the release and postsynaptic effects of DA organized at the spatiotemporal level? How does DA release at a given time affect synaptic plasticity and the transfer of information to specific neuronal subpopulations? How is dopaminergic activity regulated by various neuromodulators? Addressing these fundamental questions requires cell type–specific tools that can be used to monitor DA dynamics in real time with high spatial resolution, high molecular specificity, high sensitivity, and rapid kinetics, thereby improving our understanding of the role that DA plays in both health and disease. In this respect, our DA sensors provide a suitable tool that can be combined with other techniques such as optogenetics, calcium imaging, and electrophysiology.

Neuromodulatory neurons such as dopaminergic neurons send their projections throughout the brain, and each brain region can receive several neuromodulators. These various neurochemical pathways closely interact in order to regulate behavior in a state–dependent and highly cooperative manner. Thus, our red fluorescent rGRABDA sensors provide a new strategy for studying the interplay between various neuromodulators and neurotransmitters, given that our red fluorescent DA sensors are spectrally compatible with green fluorescent sensors such as the GRAB_ACh_ sensor^65,66^ and GRAB_NE_ sensor^67^, allowing the simultaneous monitoring of DA with acetylcholine and norepinephrine, respectively. In addition, combining the red fluorescent DA sensors with green fluorescent GCaMPs can provide new insights regarding the putative role that DA plays in regulating pre- and/or post-neuronal activities, as well as the function of target neural circuits and behaviors. Finally, several studies have shown that the GRAB–based sensor strategy can be easily used to create genetically encoded sensors based on a wide range of G-protein–coupled receptors^31,65–67^, leading to a robust and versatile multi-color toolbox that can be used to create comprehensive functional maps of neurochemical activity.

## Methods

### Animals

Postnatal 0-day-old (P0) Sprague-Dawley rats (Beijing Vital River Laboratory) and adult (P42–90) wild-type C57BL/6N (Beijing Vital River Laboratory), wild-type C57BL/6J (Charles River Laboratories), and DAT-IRES-Cre mice (Jackson Laboratory, stock number 06660) were used in this study. All animals were housed in a temperature-controlled room with a 12 h/12 h light-dark cycle. All procedures for animal surgery, maintenance, and behavior were performed using protocols that were approved by the respective animal care and use committees at Peking University, New York University, and the US National Institutes of Health.

The transgenic *Drosophila* lines UAS-rDA1m, UAS-rDA-mut, UAS-DA2m, UAS-dLight1.3b, and UAS-Pink-Flamindo were generated using Phi-C31-directed integration into attp40 or VK00005 at the Core Facility of Drosophila Resource and Technology, Shanghai Institute of Biochemistry and Cell Biology, Chinese Academy of Sciences. The following *Drosophila* lines were also used in this study: UAS-DA1m (BDSC: 80047), R13F02-Gal4 (BDSC: 49032), 30y-Gal4, and TH-Gal4-3p3-RFP^68^ (all gifts from Yi Rao, Peking University, Beijing); MB312C-Gal4 (Fly light: 2135360); MB315C-Gal4 (Fly light: 2135363); and UAS-CsChrimson-mCherry (a gift from Chuan Zhou, Institute of Zoology, Chinese Academy of Sciences, Beijing). The flies were raised on standard cornmeal-yeast medium at 25°C, 70% relative humidity, and a 12 h/12 h light-dark cycle. Adult female flies within 2 weeks after eclosion were used for fluorescence imaging.

### Molecular Biology

DNA fragments were generated using PCR amplification with primers (TSINGKE Biological Technology) containing 30 bp of overlap. The fragments were then assembled into plasmids using Gibson assembly^69^. All plasmid sequences were verified using Sanger sequencing (TSINGKE Biological Technology). For characterization in HEK293T cells, the genes expressing the red and green DA sensors were cloned into the pDisplay vector, with an IgK leader sequence inserted upstream of the sensor gene. The IRES-EGFP-CAAX gene (for red DA sensors) or IRES-mCherry-CAAX gene (for green DA sensors) was attached downstream of the sensor gene and was used as a membrane marker and to calibrate the fluorescence signal intensity. Site–directed mutagenesis was performed using primers containing randomized NNB codons (48 codons in total, encoding 20 possible amino acids) or defined codons on the target sites. For characterization in cultured neurons, the sensor genes were cloned into the pAAV vector under the control of the human synapsin promoter (hSyn). To generate stable cell lines expressing wild-type D_2_R, rDA1h, or DA2h, we generated a vector called pPacific, containing various elements, including 30 TR, the myc tag gene, a 2A sequence, the mCherry gene, the puromycin gene, and 50 TR; the genes were then cloned into the pPacific vector using Gibson assembly. Two mutations (S103P and S509G) were introduced in pCS7-PiggyBAC (ViewSolid Biotech) to generate a hyperactive piggyBac transposase^70^ for generating stable cell lines. For the TANGO assay, the wildtype D_2_R, rDA1h, or DA2h genes were cloned into the pTango vector^47^. For the luciferase complementation assay, we replaced the β_2_AR gene in the β_2_AR-Smbit construct^46^ with the wildtype D_2_R, rDA1h, or DA2h genes. To generate transgenic fly lines, the respective sensor genes were cloned into the pUAST vector, which was then used for P-element-mediated random insertion.

### Preparation and fluorescence imaging of cultured cells

HEK293T cells were cultured in DMEM (Gibco) supplemented with 10% (v/v) fetal bovine serum (Gibco) and 1% penicillin-streptomycin (Gibco) at 37°C in 5% CO_2_. The cells were plated on 96-well plates or 12 mm glass coverslips in 24-well plates and grown to 60% confluence for transfection. For transfection, the cells were incubated in a mixture containing 1 μg DNA and 3 μg PEI for 6 h. Fluorescence imaging was performed 24–48 h after transfection. Rat cortical neurons were prepared from P0 Sprague-Dawley rat pups (Beijing Vital River Laboratory). In brief, cortical neurons were dissociated from dissected rat brains in 0.25% Trypsin-EDTA (Gibco), plated on 12 mm glass coverslips coated with poly-D-lysine (Sigma-Aldrich), and cultured in Neurobasal medium (Gibco) containing 2% B-27 supplement (Gibco), 1% GlutaMAX (Gibco), and 1% penicillin-streptomycin (Gibco) at 37°C in 5% CO_2_. The neurons were transfected with an AAV expressing rDA1m, rDA1h, DA2m, DA2h, or dLight1.1 (Vigene Biosciences) after 7–9 days in culture, and fluorescence imaging was performed 3-7 days after transfection.

Cultured cells were imaged using an inverted Ti-E A1 confocal microscope (Nikon) and the Opera Phenix high-content screening system (PerkinElmer). The confocal microscope was equipped with a 40×/1.35 NA oil-immersion objective, a 488 nm laser, and a 561 nm laser. During fluorescence imaging, the cells were either bathed or perfused in a chamber containing Tyrode’s solution consisting of (in mM): 150 NaCl, 4 KCl, 2 MgCl_2_, 2 CaCl_2_, 10 HEPES, and 10 glucose (pH 7.4). Solutions containing various concentrations of drugs of DA (Sigma-Aldrich) and/or 1 μM Halo (Tocris), SCH-23390 (Tocris), Etic (Tocris), L-DOPA (Abcam), 5-HT (Tocris), histamine (Tocris), Glu (Sigma-Aldrich), GABA (Tocris), Ado (Tocris), ACh (Solarbio), NE (Tocris), Tyr (Sigma-Aldrich), or Oct (Tocris) were delivered via a custom-made perfusion system or via bath application. Between experiments, the chamber was thoroughly cleaned with 75% ethanol, 3% hydrogen peroxide, and Tyrode’s solution. GFP fluorescence was collected using a 525/50 nm emission filter, and RFP fluorescence was collected using a 595/50 nm emission filter. Photostability was measured under 1-photon illumination (confocal microscopy) using a 488 nm laser at 350 μW, and photostability was measured under 2-photon illumination using a 920 nm laser at 27.5 mW. Photobleaching was applied to the entire sensor-expressing HEK293T cell at an area of 200 μm^2^. Blue light–mediated photoactivation was measured using a 488 nm laser at 350 μW. The Opera Phenix high-content screening system was equipped with a 60×/1.15 NA water-immersion objective, a 488 nm laser, and a 561 nm laser. GFP fluorescence was collected using a 525/50 nm emission filter, and RFP fluorescence was collected using a 600/30 nm emission filter. Where indicated, the culture medium was replaced with 100 μl Tyrode’s solution containing various concentrations of the indicated drugs. The red and green sensors’ fluorescence intensity was calibrated using EGFP and mCherry, respectively.

### Spectra measurements

HEK293T cells stably expressing rDA1m, rDA1h, or DA2h were harvested and transferred to a 96-well plate in the absence or presence of 100 μM DA, and excitation and emission spectra were measured at 5 nm increments using a Safire2 multi-mode plate reader (Tecan).

### Luciferase complementation assay

The luciferase complementation assay was performed as previously described^46^. In brief, 24–48 h after transfection, HEK293T cells expressing rDA1h or DA2h were washed in PBS, harvested by trituration, and transferred to 96-well plates. DA at various concentrations (ranging from 1 nM to 100 μM) was applied to the cells, and furimazine (NanoLuc Luciferase Assay, Promega) was then applied to a final concentration of 5 μM, after which luminescence was measured using a Victor X5 multi-label plate reader (PerkinElmer).

### TANGO assay

DA was applied at various concentrations (ranging from 0.1 nM to 10 μM) to a reporter cell line stably expressing a tTA-dependent luciferase reporter and a β-arrestin2-TEV fusion gene^47^ transfected to express wild-type D_2_R, rDA1h, or DA2h. The cells were then cultured for 12 h to allow for luciferase expression. Bright-Glo (Fluc Luciferase Assay System, Promega) was then applied to a final concentration of 5 μM, and luminescence was measured using a VICTOR X5 multi-label plate reader (PerkinElmer).

### Preparation and fluorescence imaging of acute brain slices

Wild-type adult (P42-56) C57BL/6N mice were anesthetized with an intraperitoneal injection of 2,2,2-tribromoethanol (Avertin, 500 mg/kg body weight, Sigma-Aldrich), and then placed in a stereotaxic frame for AAV injection using a microsyringe pump (Nanoliter 2000 Injector, WPI). In Fig. 5a-e, AAVs expressing hSyn-rDA1m, hSyn-rDA1h, or hSyn-DA2m (Vigene Biosciences) were injected (400 nl per injection site) into the NAc using the following coordinates: AP: +1.4 mm relative to Bregma, ML: ±1.2 mm relative to Bregma, depth: 4.0 mm from the dura. In Fig. 5f-k, the AAV expressing hSyn-rDA1m (Vigene Biosciences) was injected (400 nl per injection site) into the NAc using the coordinates listed above, and the AAV expressing hSyn-axon-GCaMP6s (BrainVTA) was injected (400 nl per injection site) into the VTA using the following coordinates: AP: −3.2 mm relative to Bregma, ML: ±0.5 mm relative to Bregma, depth: 4.1 mm from the dura.

Two weeks after virus injection, the mice were anesthetized with an intraperitoneal injection of Avertin (500 mg/kg body weight) and perfused with ice-cold oxygenated slicing buffer containing (in mM): 110 choline-Cl, 2.5 KCl, 1 NaH_2_PO_4_, 25 NaHCO_3_, 7 MgCl_2_, 25 glucose, and 0.5 CaCl_2_. The brains were immediately removed and placed in ice-cold oxygenated slicing buffer. The brains were sectioned into 300 μm thick slices using a VT1200 vibratome (Leica), and the slices were incubated at 34°C for at least 40 min in oxygenated artificial cerebrospinal fluid (ACSF) containing (in mM): 125 NaCl, 2.5 KCl, 1 NaH_2_PO_4_, 25 NaHCO_3_, 1.3 MgCl_2_, 25 glucose, and 2 CaCl_2_. For fluorescence imaging, the slices were transferred to an imaging chamber and placed under an FV1000MPE 2-photon microscope (Olympus) equipped with a 25×/1.05 NA water-immersion objective and a mode–locked Mai Tai Ti: Sapphire laser (Spectra-Physics). A 950 nm laser was used to excite rDA1m and rDA1h, and fluorescence was collected using a 575–630 nm filter. A 920 nm laser was used to excite DA2m, and fluorescence was collected using a 495–540 nm filter. For electrical stimulation, a bipolar electrode (cat. number WE30031.0A3, MicroProbes for Life Science) was positioned near the core of the NAc using fluorescence guidance. Fluorescence imaging and electrical stimulation were synchronized using an Arduino board with custom-written programs. All images collected during electrical stimulation were recorded at a frame rate of 0.1482 s/frame with 128×96 pixels per frame. The stimulation voltage was 4–6 V, and the duration of each stimulus was 1 ms. Drugs were applied to the imaging chamber by perfusion at a flow rate at 4 ml/min.

### Fluorescence imaging of transgenic flies

Adult female flies (within 2 weeks after eclosion) were used for fluorescence imaging. For imaging, the fly was mounted to a customized chamber using tape such that the antennae and abdomen were exposed to the air. A section of rectangular cuticle between the eyes, as well as the air sacs and fat bodies, were then removed to expose the brain, which was bathed in adult hemolymph-like solution (AHLS) containing (in mM): 108 NaCl, 5 KCl, 5 HEPES, 5 trehalose, 5 sucrose, 26 NaHCO_3_, 1 NaH_2_PO_4_, 2 CaCl_2_, and 1-2 MgCl_2_. Fluorescence imaging was performed using an FV1000MPE 2-photon microscope (Olympus) equipped with a 25×/1.05 NA water-immersion objective and a mode–locked Mai Tai Ti: Sapphire laser (Spectra-Physics). A 950 nm laser was used to excite rDA1m and Pink-Flamindo, and a 575-630 nm filter was used to collect the fluorescence signals. A 930 nm laser was used to excite DA2m and dLight1.3b, and a 495-540 nm filter was used to collect the fluorescence signals. For odorant stimulation, isoamyl acetate (Sigma-Aldrich) was first diluted 200-fold in mineral oil in a bottle and then diluted 5-fold in air, which was then delivered to the fly’s antennae at a rate of 1000 ml/min. Halo (Tocris) was added directly to the AHLS at the final concentration. For electrical stimulation, a glass electrode (0.2 MΩ resistance) was placed in the olfactory mushroom body in the vicinity of dopaminergic neurons, and the stimulation voltage was set at 20–80 V. For body shock, two wires were attached to the fly’s abdomen, and 60 V electrical pulses (500 ms duration) were delivered. For DA perfusion, a patch of the blood-brain barrier was carefully removed using tweezers, and AHLS containing 100 μM DA was applied to exchange the normal bath solution. For optogenetic stimulation, a 200 mW 635 nm laser (Changchun Liangli photoelectricity Ltd.) was used to deliver light to the fly brain via an optical fiber. An Arduino board with custom–written programs was used to synchronize the stimulation and fluorescence imaging. The sampling rate during odorant stimulation, electrical stimulation, body shock, and DA perfusion was 6.7 Hz, 12 Hz, 6.7 Hz, and 1 Hz, respectively.

### Fiber photometry recording of optogenetically induced DA release in freely moving mice

Adult (P42–56) DAT-IRES-Cre mice (Jackson Laboratory, stock number 06660) were anesthetized with isoflurane and placed in a stereotaxic frame for AAV injection. AAVs expressing hSyn-rDA1m, hSyn-rDA1h, hSyn-rDA-mut, hSyn-DA2m, or hSyn-DA2h (Vigene Biosciences) as well as hSyn-EGFP (Addgene, cat. number 50465) or hSyn-tdTomato (a gift from Dr. Yakel’s Lab) were injected (1 μl per site) into the dorsal striatum using the following coordinates: AP: −0.5 mm relative to Bregma, ML: ±2.4 mm relative to Bregma, depth: 2.2 mm from the dura. The AAV expressing Ef1α-DIO-C1V1-YFP (NIEHS Viral Vector Core) was injected (500 nl per site) into the SNc using the following coordinates: AP: −3.1 mm relative to Bregma, ML: ±1.5 mm relative to Bregma, depth: 4.0 mm from the dura. Optical fibers (105 μm core/125 μm cladding) were implanted in the dorsal striatum and SNc 4 weeks after AAV injection. Fiber photometry recording in the dorsal striatum was performed using a 488 nm laser at 50 μW for DA2m and DA2h, a 488 nm laser at 1 μW and a 561 nm laser at 50 μW for rDA1m, rDA1h, and rDA-mut. C1V1 in the SNc was stimulated using a 561 nm laser at 9.9 mW. The measured emission spectra were fitted using a linear unmixing algorithm (https://www.niehs.nih.gov/research/atniehs/labs/ln/pi/iv/tools/index.cfm). To evoke C1V1-mediated DA release, pulse trains (10 ms pulses at 20 Hz for 1 s) were delivered to the SNc using a 561 nm laser at 9.9 mW. To avoid signal decay, the excitation lasers were controlled using an optical shutter (Thorlabs) in which the shutter was turned on 10 s before the 561 nm pulse trains and turned off 35 s after stimulation.

### Fiber photometry recording of DA dynamics in the NAc during sexual behavior

Adult (P60–90) wild-type male C57BL/6J mice (Charles River Laboratories) were anesthetized with isoflurane and placed in a stereotaxic frame for AAV injection. AAVs expressing hSyn-rDA1m or hSyn-DA2h (Vigene Biosciences) were injected (80–140 nl per site) into the NAc using the following coordinates: AP: +0.98 mm relative to Bregma, ML: ±1.2 mm relative to Bregma, depth: 4.6 mm from the dura. For the co-expression of rDA1m and DA2h in Fig. 6g-l, AAVs expressing hSyn-rDA1m and hSyn-DA2h were injected at a 1:1 ratio. After AAV injection, optical fibers (400 μm diameter) were implanted in the NAc, and fiber photometry recording was performed two weeks after AAV injection. The setups for bilateral recording and dual-color recording are shown in Fig. 6a and Fig. 6g, respectively. In brief, a 311 Hz 472/30 nm filtered LED (Thorlabs) at 30 μW was used to excite DA1h and DA2h, and a 400 Hz 590/20 nm filtered LED (Thorlabs) at 30 μW was used to excite rDA1m. A 535/50 nm filter was used to collect the fluorescence signal from DA1h and DA2h, and a 524/628-25 nm dual-band bandpass filter was used to collected the fluorescence signal from rDA1m and DA2h during the dual-color recording. The various sexual behaviors are defined as previously described^31^ following published conventions^71^. For immunofluorescence, the mice were anesthetized and then perfused with 4% paraformaldehyde (PFA). The brains were removed, fixed in 4% PFA for 4 h, and then cryoprotected in 20% (w/v) sucrose for 24 h. The brains were then embedded in tissue-freezing medium and sectioned into 60 μm thick slices using a CM1900 cryostat (Leica). rDA1m was immunostained using a rabbit anti-RFP antibody (1:1000, Takara, cat. number 632496) followed by a Cy3-conjugated donkey anti-rabbit secondary antibody (1:1000, Jackson ImmunoResearch, cat. number 113713). DA1h and DA2h were immunostained using a chicken anti-GFP antibody (1:1000, Abcam, cat. number ab13970) followed by an Alexa 488-conjugated donkey anti-chicken secondary antibody (1:1000, Jackson ImmunoResearch, cat. number 116967).

### Quantification and statistical analysis

Imaging data from cultured cells, acute brain slices, and transgenic flies were processed using ImageJ software (NIH). Pseudocolor images were generated using custom–written programs. The fluorescence response (ΔF/F_0_) was calculated using the formula [(F-F_0_)/F_0_], in which F_0_ is the baseline fluorescence signal. The signal-to-noise ratio (SNR) was calculated as the peak response divided by the standard error of the baseline fluorescence fluctuation. Summary data are presented as the mean ± s.e.m., and group data were analyzed using the Student’s *t*-test, one-way ANOVA (Supplementary Fig. S5) or two-way ANOVA (Fig. 6). Where indicated, \**p* < 0.05, \*\**p* < 0.01, \*\*\**p* < 0.001, and n.s., not significant *(p* > 0.05).

### Data and software availability

Plasmids expressing the sensors used in this study were deposited at Addgene (https://www.addgene.org/Yulong_Li/). The custom–written programs will be provided upon request to the lead contact, Yulong Li.

## Acknowledgments

This work was supported by the Beijing Municipal Science & Technology Commission (Z181100001318002), the Beijing Brain Initiative of Beijing Municipal Science & Technology Commission (Z181100001518004), Guangdong Grant ‘Key Technologies for Treatment of Brain Disorders’ (2018B030332001), the General Program of National Natural Science Foundation of China (projects 31671118, 31871087, and 31925017), the NIH BRAIN Initiative (NS103558), and grants from the Peking-Tsinghua Center for Life Sciences and the State Key Laboratory of Membrane Biology at Peking University School of Life Sciences to Y.L.; the NIH (grants R01MH101377 and R21HD090563) and an Irma T. Hirschl Career Scientist Award to D.L.; and the Intramural Research Program of the US NIH/NIEHS (1ZIAES103310) to G.C.. We thank Yi Rao for sharing the 2-photon microscope and Xiaoguang Lei at PKU-CLS for providing support for the Opera Phenix high-content screening system.

## Author contributions

Y.L. supervised the study. F.S., Y.L. designed the study. F.S., Y. Zhuo, and Y. Zhang performed the experiments related to developing, optimizing, and characterizing the sensors in cultured HEK293T cells and neurons with help from J.F., H.D. and C.Q.. F.S. and T.Q. performed the surgery and 2-photon imaging experiments related to the validation of the sensors in acute brain slices. X.L., K.T., and J. Zeng performed the 2-photon imaging experiments in transgenic flies. J. Zhou performed the fiber photometry recordings during optogenetics in freely moving mice under the supervision of G.C.. B.D. performed the fiber photometry recordings in the mouse NAc during sexual behavior under the supervision of D.L.. All authors contributed to the data interpretation and analysis. F.S. and Y.L. wrote the manuscript with input from all authors.

## Competing interests

F.S. and Y. L. have filed patent applications whose value might be affected by this publication.

---

**Supplementary Fig. S1.**
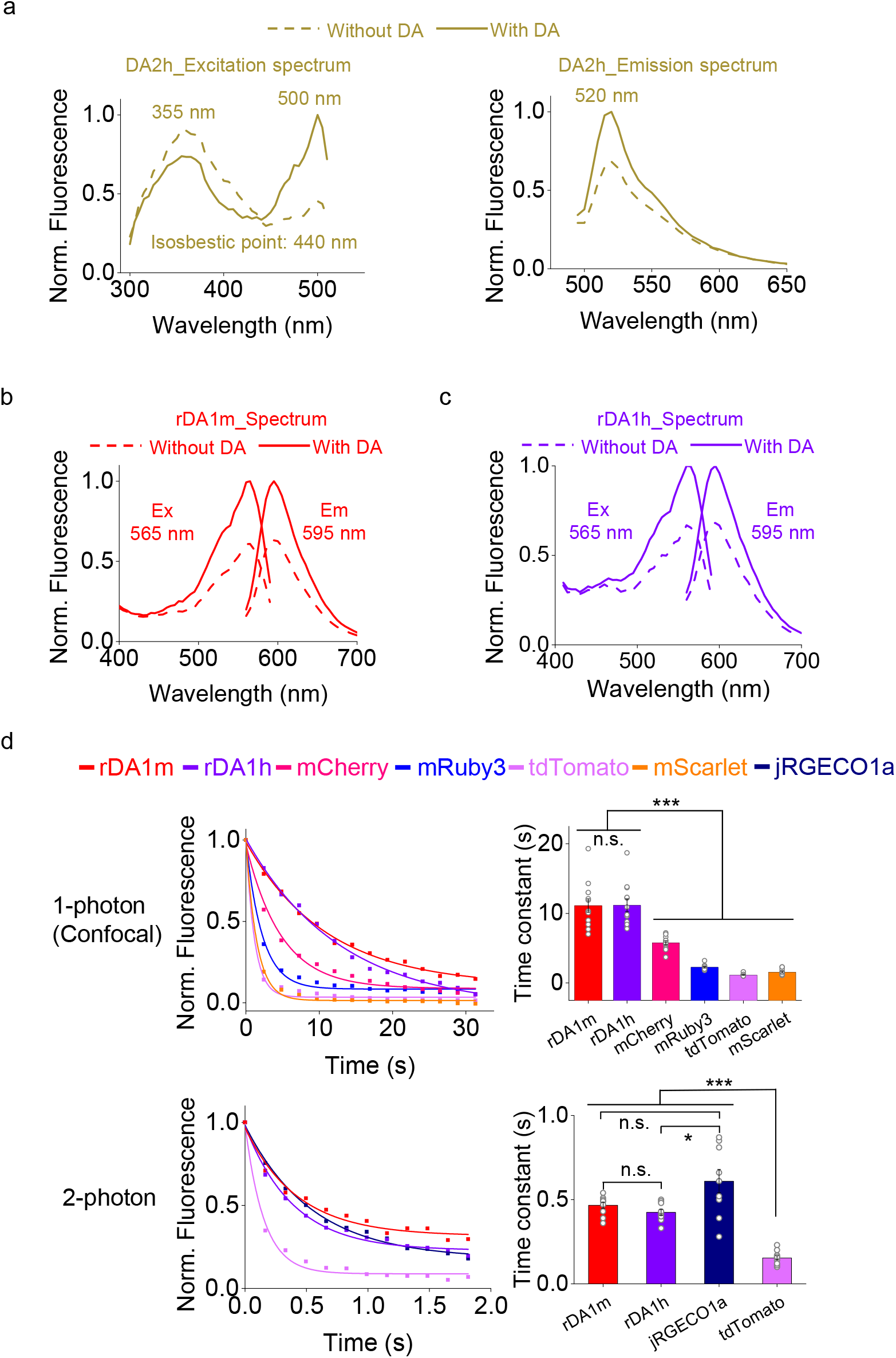
Characterization of the sensors expressed in HEK293T cells (related to Fig. 2). **a-c**, The excitation and emission spectra of DA2h (**a**), rDA1m (**b**), and rDA1h (**c**) in the absence and presence of DA. **d**, The photostability of rDA1m and rDA1h (in the presence of 100 μM DA) and the indicated fluorescent proteins was measured using 1-photon (top) and 2-photon (bottom) microscopy. Left, representative photobleaching curves; each curve was fitted to a single-exponential function. Right, group summary of the photobleaching time constants (n=10–12 cells each).

**Supplementary Fig. S2.**
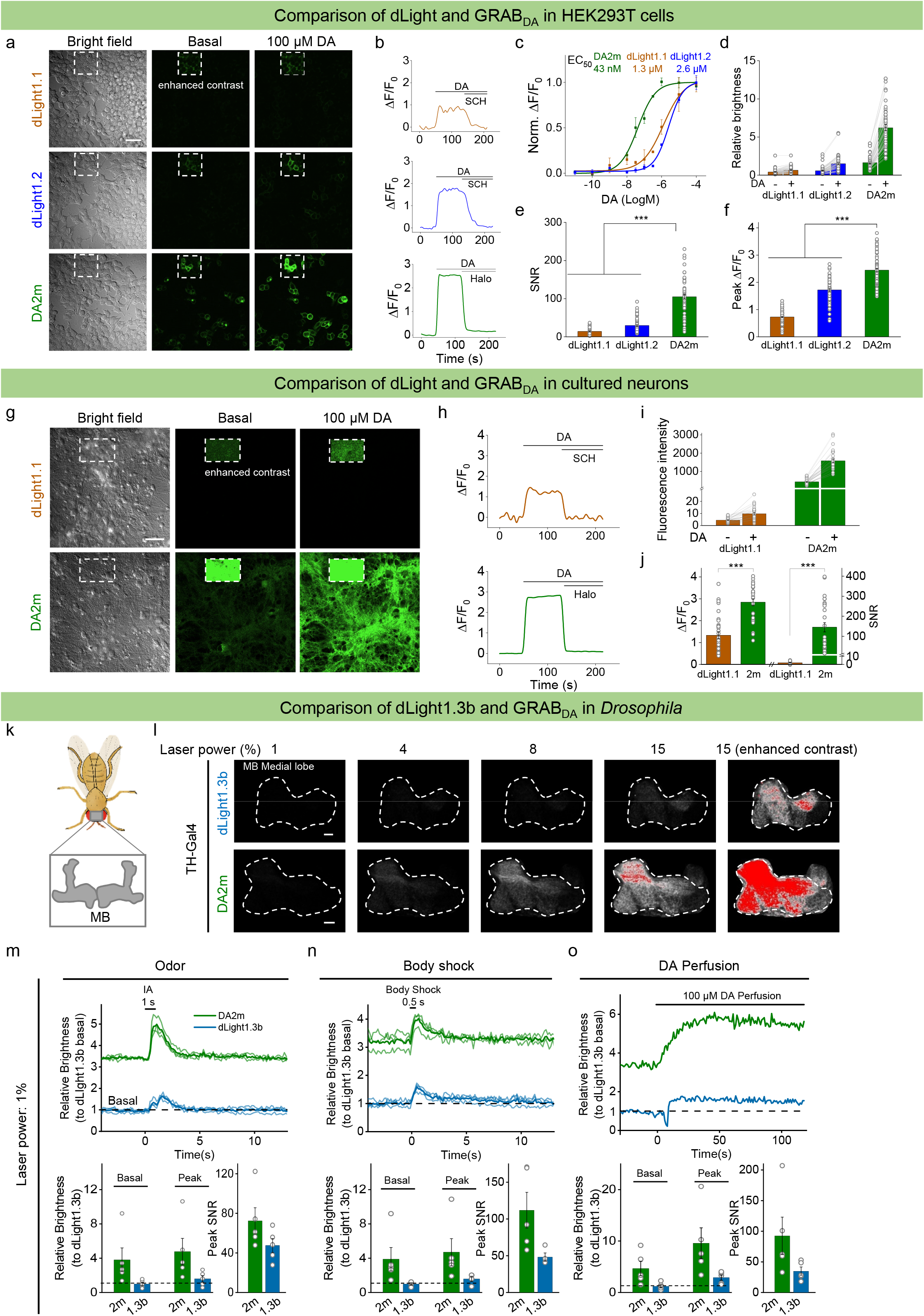
Comparison between dLight and GRAB_DA_ (related to Figs. 2 and 4). **a**, Representative bright-field and fluorescence images measured before (baseline) and after application of DA in HEK293T cells expressing dLight1.1, dLight1.2, or DA2m. Scale bar, 50 μm. **b**, Representative traces of dLight1.1, dLight1.2, and DA2m ΔF/F_0_ in response to 100 μM DA followed by either 10 μM SCH (dLight1.1 and dLight1.2) or 10 μM Halo (DA2m). **c**, Normalized dose-response curves for dLight1.1, dLight1.2, and DA2m (n=3 wells with 100–500 cells/well). **d-f**, Group summary of the relative brightness (**d**), signal-to-noise ratio (SNR) (**e**), and peak ΔF/F_0_ (**f**) for dLight1.1, dLight1.2, and DA2m in response to 100 μM DA (n=35–77 cells each). To assess brightness, the membrane–targeted red fluorescent protein mCherry-CAAX was co-expressed with each sensor, and relative brightness was normalized to the corresponding mCherry signal. **g-j**, Similar to **a-f**, except that dLight1.1 and DA2m were expressed in cultured neurons (n=28–30 neurons each). Scale bar, 50 μm. **k**, Schematic illustration depicting the location of the *Drosophila* olfactory mushroom body (MB). **l**, Fluorescence images of the MB in flies expressing dLight1.3b (top) or DA2m (bottom) using 2-photon microscopy at the indicated laser power settings. Enhanced-contrast images at 15% laser power are shown at the right. Fluorescence is shown in grayscale, with saturated pixels shown in red. Scale bars, 10 μm. **m-o**, Representative traces (top) and group summary of relative brightness measured for dLight1.3b and DA2m during odorant application (**m**), body shock (**n**), and DA perfusion (**o**); n=4–5 flies each. Average traces (bold) overlaid with single-trial traces (light) from one fly are shown for representation.

**Supplementary Fig. S3.**
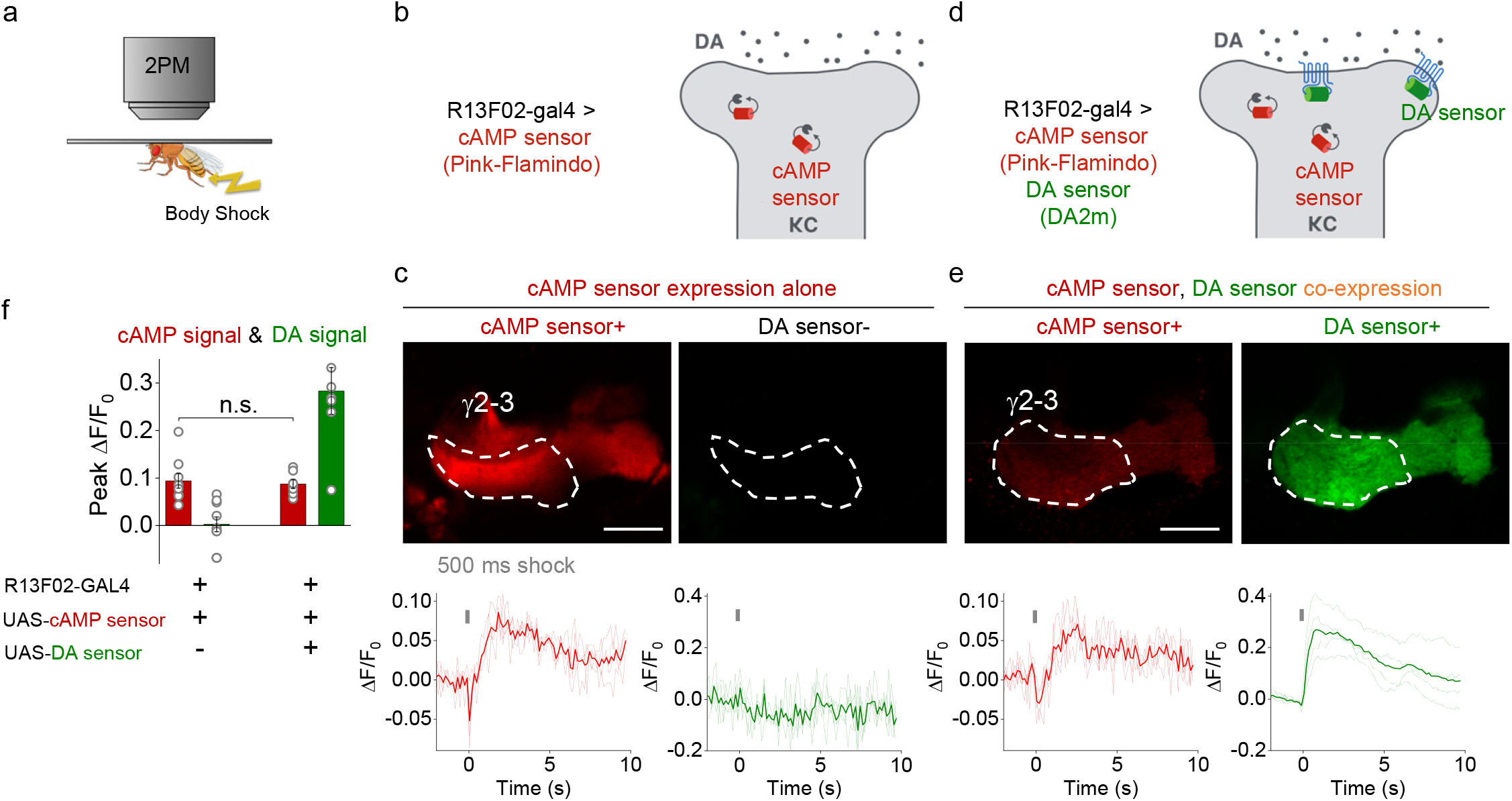
Expressing GRABDA2m sensors does not affect cAMP signaling *in vivo* (related to Fig. 4). **a**, Schematic illustration depicting the experimental setup for imaging body shock–evoked DA release using 2-photon microscopy. **b-e**, Schematic illustrations depicting the experimental strategy (**b,d**) and representative fluorescence images and ΔF/F_0_ traces (**c,e**) in flies expressing the cAMP sensor Pink-Flamindo (**b,c**) or co-expressing Pink-Flamindo and DA2m (**d,e**) in MB KCs. Where indicated, a 500 ms body shock was delivered. The ROIs for measuring the γ2-γ3 compartments in the MB are indicated by dashed white lines. Scale bars, 25 μm. Average traces (bold) overlaid with single-trial traces (light) from one fly are shown for representation. **f**, Group summary of peak Pink-Flamindo and DA2m ΔF/F_0_ measured under the indicated conditions (n=7–9 flies each).

**Supplementary Fig. S4.**
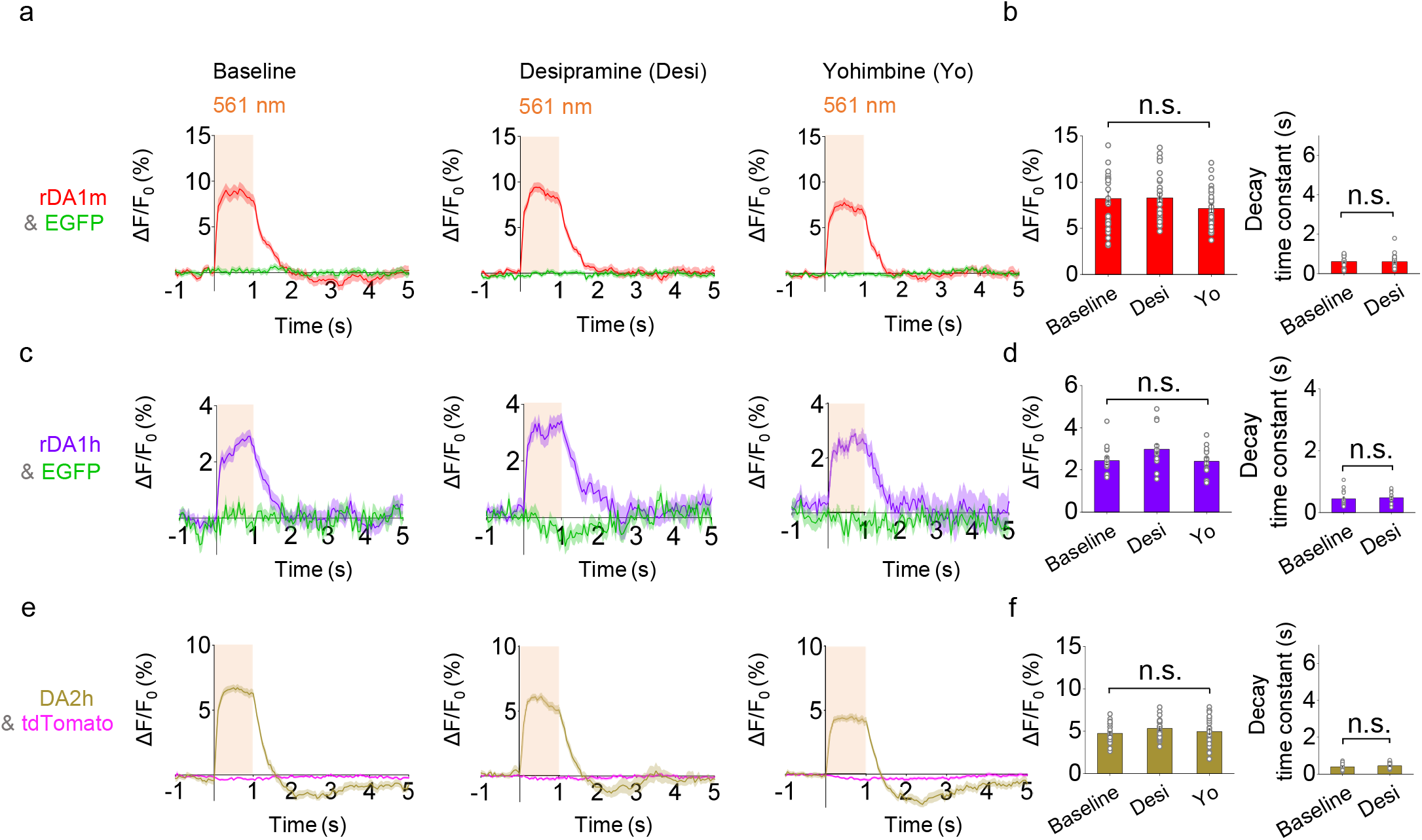
Optogenetically induced nigrostriatal DA release in freely moving mice is not affected by desipramine or yohimbine (related to Fig. 5). **a, c,e**, Average traces of ΔF/F_0_ in mice expressing rDA1m and EGFP (**a**), rDA1h and EGFP (**c**), or DA2h and tdTomato (**e**) in the dorsal striatum. Where indicated, the experiments were conducted in mice treated with either the norepinephrine transporter blocker desipramine or the α2AR antagonist yohimbine. **b, d,f**, Group summary of ΔF/F_0_ and τ_off_ for the experiments shown in **a, c**, and **e**, respectively (n=25–30 trials from 5-6 hemispheres of 3-6 mice each).

**Supplementary Fig. S5.**
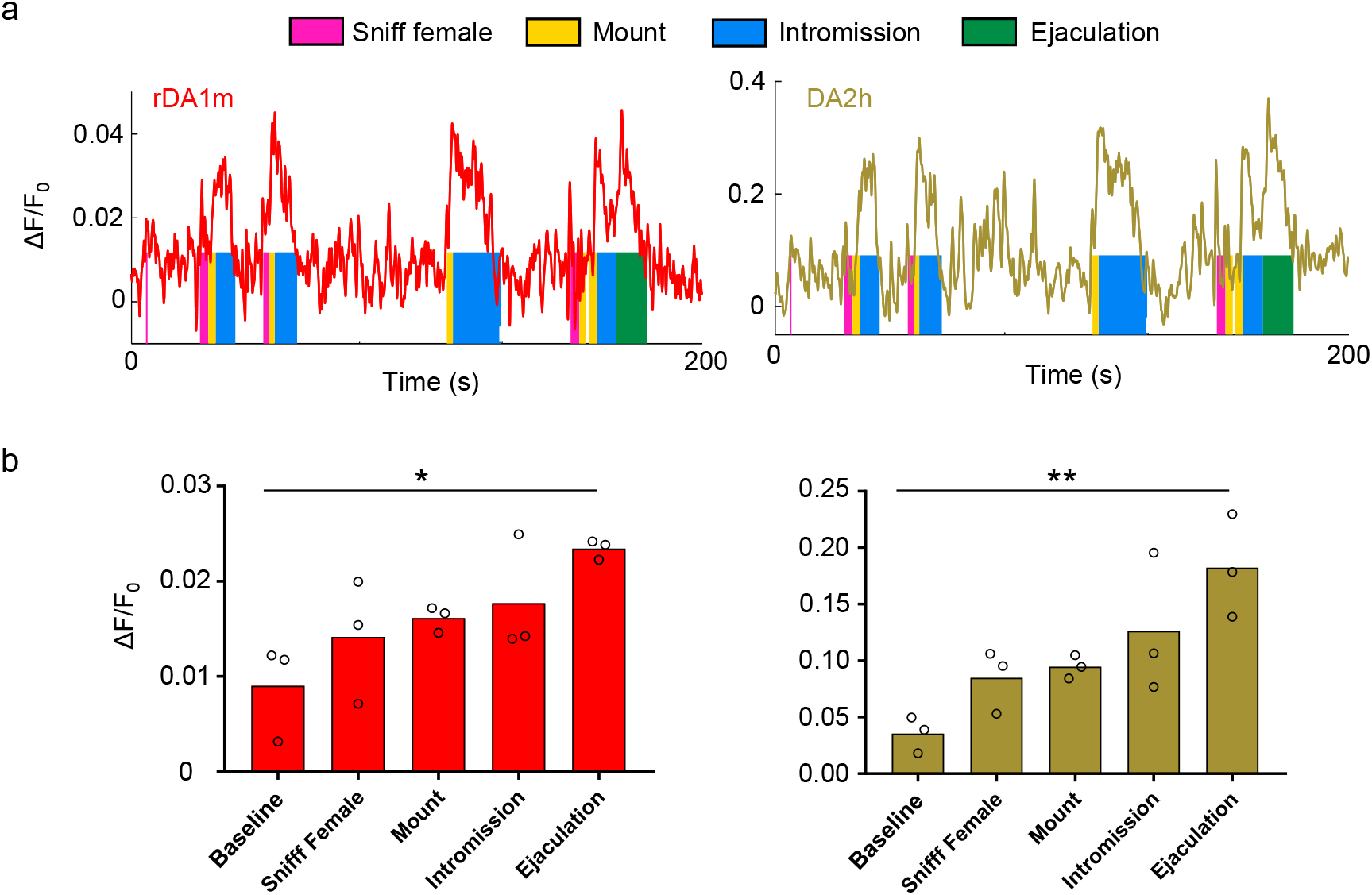
The ΔF/F_0_ signals of rDA1m and DA2h during sexual behavior (related to Fig. 6). **a**, Representative traces of rDA1m and DA2h ΔF/F_0_ measured during the indicated stages of mating. **b**, Group summary of the ΔF/F_0_ measured for rDA1m and DA2h during the indicated mating events (n=3 mice).

**Supplementary Fig. S6.**
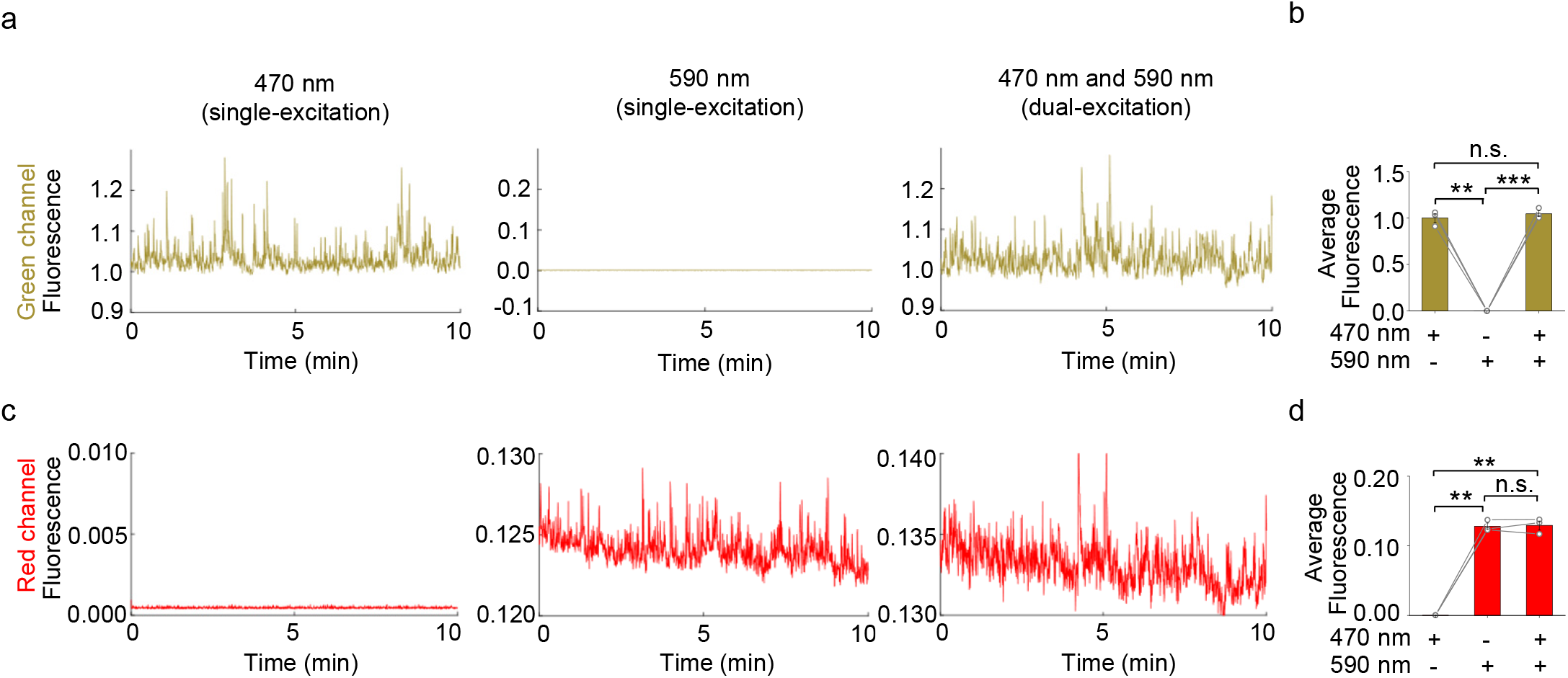
The excitation wavelength for DA2h does not excite rDA1m, and *vice versa* (related to Fig. 6). Mice co-expressing DA2h and rDA1m in the NAc were used for these experiments. Delivery of 470 nm light excited DA2h (green) but had no effect on rDA1m (red); conversely, delivery of 590 nm light excited rDA1m but had no effect on DA2h. Delivery of both 470 nm and 590 nm light using dual-wavelength excitation excited both DA2h and rDA1m. Shown at the right are the group summaries (n=3 mice each).

## Notes

#### Summary of Updates

The distribution/reuse options are changed.

